# gmxapi: a GROMACS-native Python interface for molecular dynamics with ensemble and plugin support

**DOI:** 10.1101/2021.07.18.452496

**Authors:** M. Eric Irrgang, Caroline Davis, Peter M. Kasson

## Abstract

Gmxapi provides an integrated, native Python API for both standard and advanced molecular dynamics simulations in GROMACS. The Python interface permits multiple levels of integration with the core GROMACS libraries, and legacy support is provided via an interface that mimics the command-line syntax, so that all GROMACS commands are fully available. Gmxapi has been officially supported since the GROMACS 2019 release and is enabled by default in current versions of the software. Here we describe gmxapi 0.3 and later. Beyond simply wrapping GROMACS library operations, the API permits several advanced operations that are not feasible using the prior command-line interface. First, the API allows custom user plugin code within the molecular dynamics force calculations, so users can execute custom algorithms without modifying the GROMACS source. Second, the Python interface allows tasks to be dynamically defined, so high-level algorithms for molecular dynamics simulation and analysis can be coordinated with loop and conditional operations. Gmxapi makes GROMACS more accessible to custom Python scripting while also providing support for high-level data-flow simulation algorithms that were previously feasible only in external packages.

**Author Summary:** The gmxapi software provides a Python interface for molecular dynamics simulations in GROMACS. In addition to simply wrapping GROMACS commands, it supports custom user plugin code, ensemble simulation, and data-flow chaining of commands. As such, gmxapi enables the writing and execution of high-level simulation algorithms. The software ships with GROMACS and is freely available under an LGPL2 license.

## Introduction

As molecular dynamics simulations have become more complex and mature as scientific tools, typical simulation use is shifting from manual invocation of a few simulations and analysis tools to pre-defined simulation and analysis protocols, often involving many simulation trajectories. In addition, custom applications for advanced sampling[1-4] or molecular structure refinement[5-7] are becoming more common, where the molecular dynamics engine is used as part of a more complex data integration protocol. In these more complex use scenarios, toolchain and file-system management as well as execution of many simulations can become limiting factors. As a result, many scientists either develop custom scripts, custom modifications of the simulation source code, or adapt general-purpose workflow engines[8-10] to molecular simulation tasks. There also exist some molecular-simulation-specific workflow engines, frequently coupled to the underlying molecular simulation code via command-line interfaces[11-16]. All of the above can be brittle, particularly without robust APIs for simulation interfaces to workflow engines. Integration of parallel analysis into unified jobs can also be challenging whether this is performed via built-in tool parallelism or user-level coding, such as Python multiprocessing packages or workflow/execution managers[9,10,12,13,17-19].

We previously reported on gmxapi 0.0.4[20], which allows Python driven molecular dynamics in GROMACS[21,22] to be extended at run time with custom researcher code. Here, we describe features present in gmxapi 0.2 and beyond, which offers both more advanced ensemble simulation logic and a feature-complete interface for GROMACS tools and analysis. This framework allows ensemble methods to be implemented with less code, and without patching an official GROMACS release. Data-flow-oriented programming logic and general paradigm for integrating new software tools further enhance the utility of the package. Both the libgmxapi C++ interface and the gmxapi Python package are now maintained and distributed with GROMACS; gmxapi 0.2 is integrated with GROMACS 2021, and gmxapi 0.3 is integrated with GROMACS 2022.

### Design and Implementation

We outline distinguishing features and key user functionality of gmxapi, followed by a more technical design discussion. Gmxapi offers a Python scripting interface maintained as part of the GROMACS software, and thus one basic feature is the ability to reproduce all GROMACS command-line calls. This is done through a combination of native gmxapi calls such as gmxapi.mdrun() and a wrapper that permits any operation from the GROMACS command-line client: gmxapi.commandline_operation(). These are illustrated in Figure 1a. We also highlight three key additional features not possible with a simple wrapper script: built-in ensemble parallelism, composability, and plugins.

**Figure 1:**
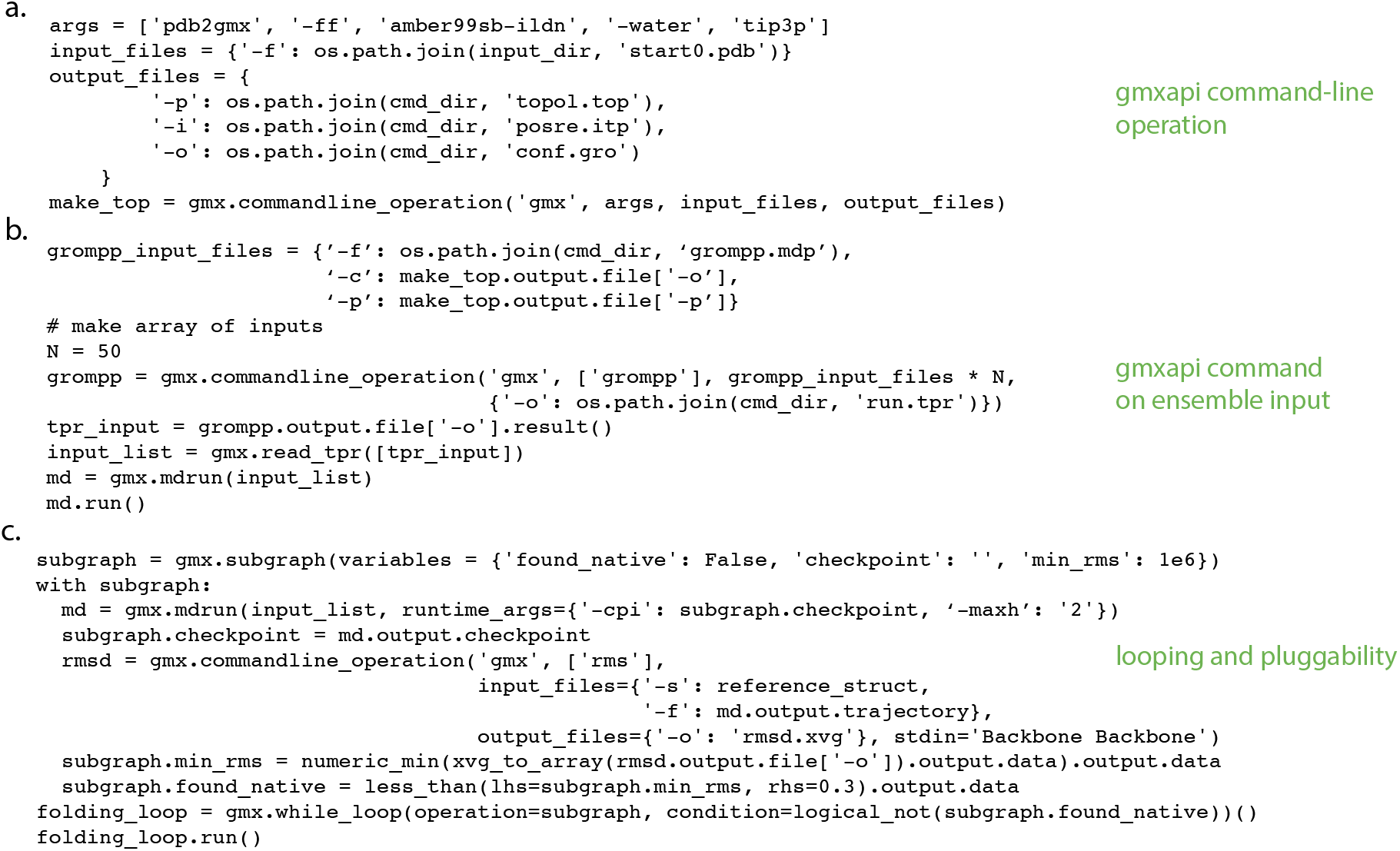
gmxapi usage examples. Panel (a) shows an example of the commandline_operation function by which gmxapi can reproduce any GROMACS functionality. Panel (b) shows gmxapi molecular dynamics calls operating on ensemble input, providing straightforward high-level parallelism in addition to parallelization within each command. Panel (c) demonstrates both a while loop and pluggability of gmxapi components. Together, the examples will execute an ensemble of small protein folding simulations until at least one ensemble member samples the native state.

### Ensemble Parallelism

Simulations are increasingly being performed not singly but as a collection of related tasks, which we will term an ensemble. This collection could consist of a set of replicas sampling a thermodynamic ensemble, but it could also represent a parameter sweep across experimental conditions or a set of simulations with different starting conformations as part of an advanced sampling strategy. Ensembles are treated as first-class objects in gmxapi, and the python interface is designed to facilitate parallelism across such ensembles, which is in turn implemented by the gmxapi backend. This is illustrated in Figure 1b. The ability to flexibly and simply articulate high-level ensemble logic is thus a key feature of gmxapi.

### Composability

Another key design and usability feature of gmxapi is the notion of composability. Similar to Tinkertoys, different calls to GROMACS commands through the API are designed to be plugged together. This is illustrated in Figure 1c and has two advantages: a natural way to conceptualize a sequence of GROMACS calls (or calls to external programs that can use the gmxapi wrapper facility) and a way to parallelize using the ensemble logic and data-flow management by gmxapi.

### Plugins

Gmxapi includes the ability to add user-defined plugins that can interact with GROMACS at runtime without modifying the GROMACS source. Plugins are currently used to implement custom force routines; these run using the native GROMACS parallel decomposition and thus can benefit from parallelism as well as acceleration. More details are given in the technical design section below.

### Technical Design

We used pybind11[23] to implement Python bindings to a C++ API for the GROMACS[21,22] molecular simulation library. The design concept of gmxapi, as with many modern high-level interfaces, is for the user to construct a computational graph where execution is then managed by lower layers of the software stack. Similar to Keras[24], we recognize that explicit data-flow programming is not natural for all users and thus provide an interface where data-flow can be specified explicitly or implicitly. Figure 2 illustrates the gmxapi paradigm in which control signals and data are treated in the same framework. Branching and other control events require that some work is “dynamic”, such that the complete workflow graph may not be determined until runtime. However, the workflow can be fully specified in terms of workflow commands and data primitives (Figure 3).

**Figure 2:**
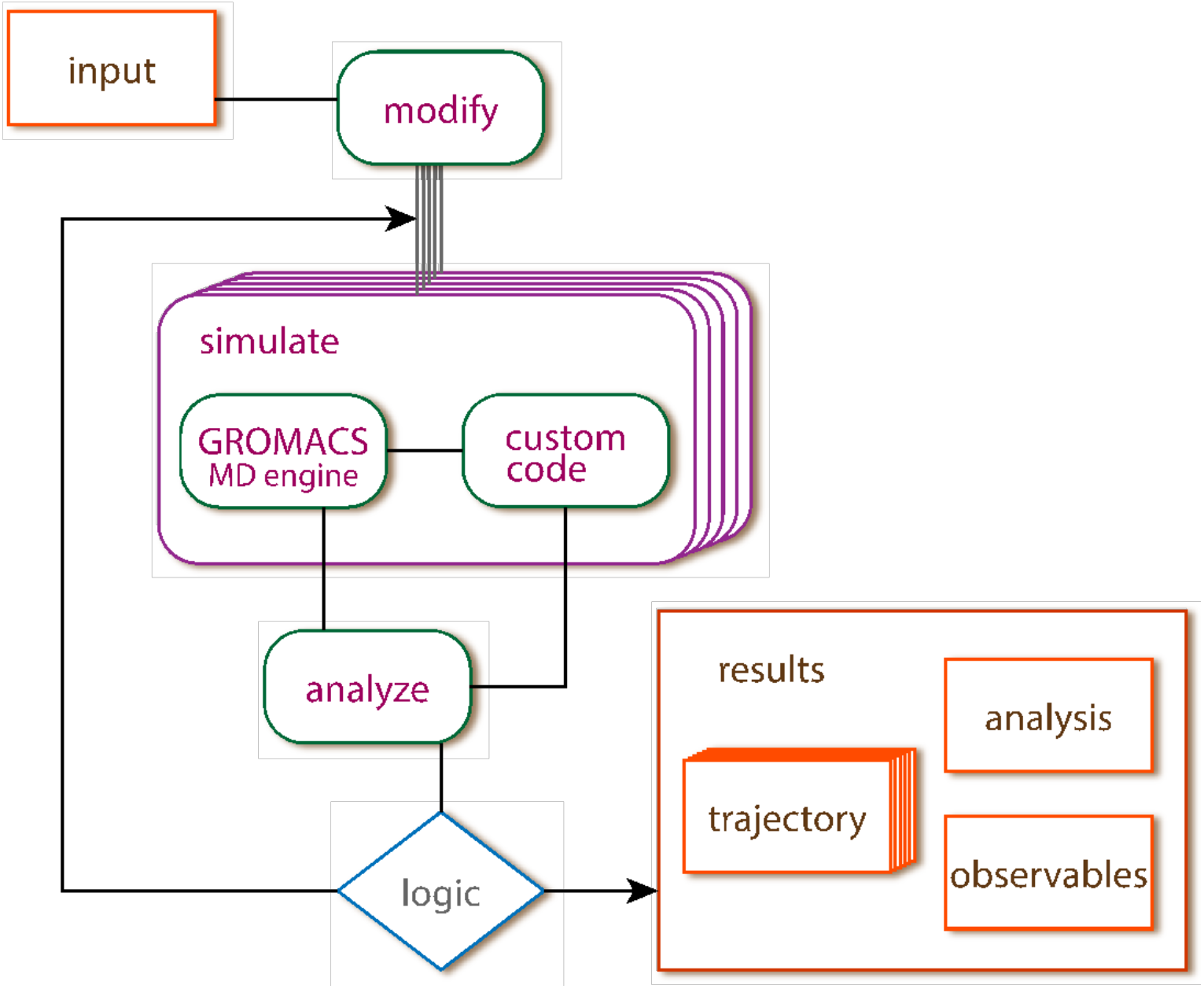
Schematic of data flow and control flow for a segment of a complex simulation workflow. Ensembles of simulations can be run (denoted by stacked rectangles) by gmxapi merely by passing an array of inputs instead of a single input. Custom plugins can interact with running MD simulations. Finally, conditional and looping logic can create high-level simulation algorithms.

**Figure 3:**
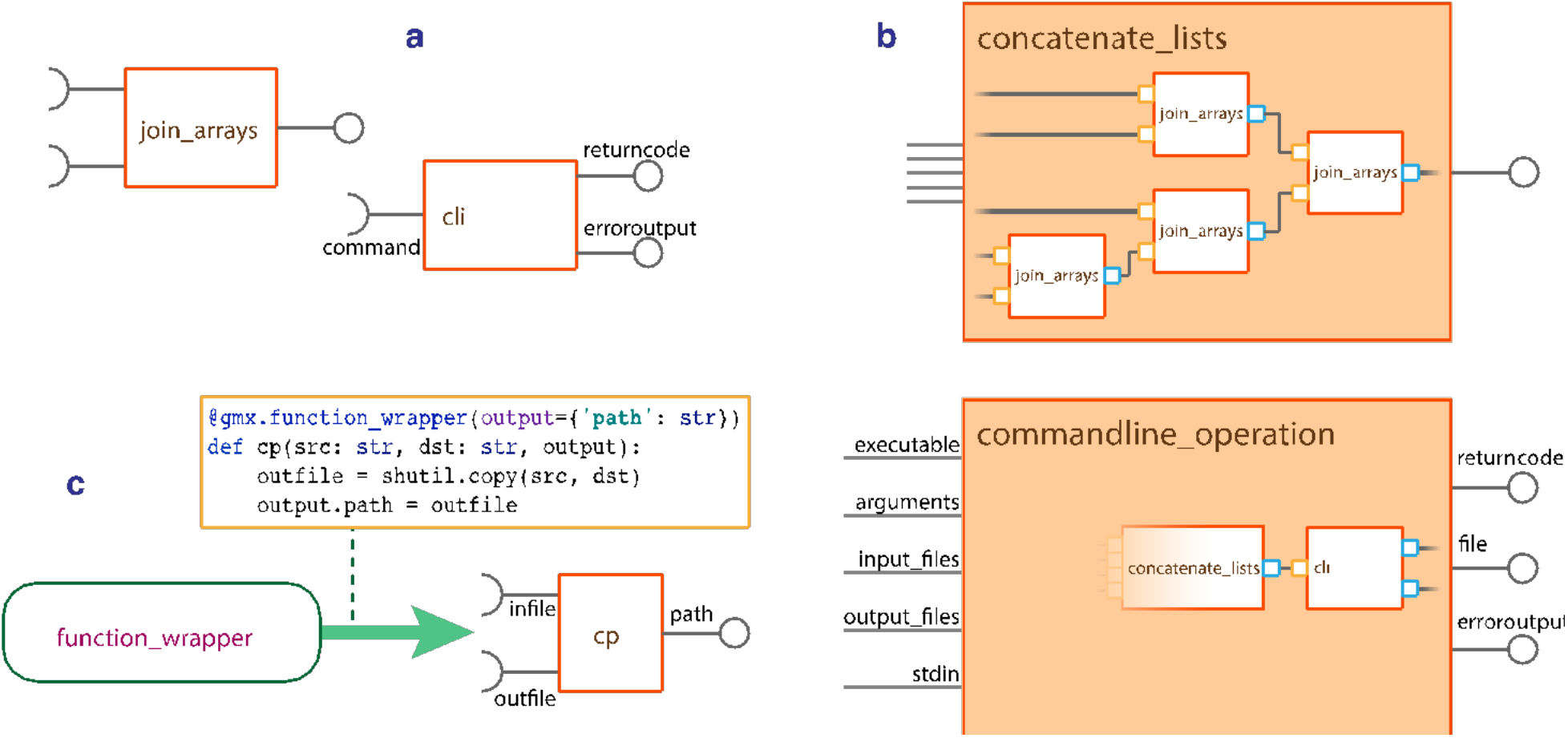
Schematics for gmxapi operations. As shown in panel (a), gmxapi operations have well defined inputs and outputs. These can operate on arrays of inputs and also include support for legacy GROMACS operations by wrapping the command-line toolset, as shown in panel (b). As shown in panel (c), the decorator @function_wrapper allows arbitrary user code to be transformed into a gmxapi operation.

### Data flow formalism

Data-flow formalisms have become increasingly popular in contemporary workflow engines due to their ability to separate work specification and task execution, permitting more straightforward resolution of dependencies and optimization of computation and data movement. For the same reasons, execution in gmxapi is deferred as much as possible to when and where it is required. The high-level gmxapi Python interface creates and executes a directed acyclic graph (DAG) specifying the computational work to be done. Nodes of the work graph represent discrete *operations* that produce and consume well defined inputs and outputs. Operation and data references in the work graph are proxy objects for the computational tasks and data until the graph is run. Execution management may be optimized by running only as much of the graph as necessary to satisfy explicit data dependencies. In addition, in order to minimize unnecessary data movement, most gmxapi operations do not transfer data from the C++ library to the Python interpreter unless explicitly requested by the user. When a gmxapi command is used to add work to the work graph, it returns a reference to the graph node representing the operation. The operation’s output attributes may be used as inputs to further commands, and may be used as “Future” results, as discussed below. Calling “result()” on a Future forces dependency resolution and data localization.

### Python interface

The gmxapi Python interface consists of five categories of operations: 1) typing and logical operations, 2) bound GROMACS API calls, 3) legacy GROMACS command-line operations, 4) utilities for user creation of new gmxapi operations, and 5) looping and conditional operations. These are described in sequence below.

The simplest gmxapi operations manipulate signals, data topology, or typing. “join_arrays”, “logical_not”, and “make_constant” have strict definitions for inputs and outputs (Figure 3a). These operations are also composable: “concatenate_lists” is a helper function that reduces a number of inputs in terms of “join_arrays” (Figure 3b).

The preferred mechanism for interfacing to GROMACS is via bound API calls. Commands like “read_tpr”, “modify_input”, and “mdrun” use a binary Python extension module written in C++ using pybind11 to interact with GROMACS operations and data through libgmxapi, which is a C++ library installed by default with recent versions of GROMACS.

In order to provide legacy support for the full range of GROMACS command-line operations, “gmxapi.commandline.cli()” is a simple pure-Python gmxapi operation that wraps command-line tools. It is a thin wrapper of the Python “subprocess” module, so command inputs and outputs are embedded in the argument list parameter. Instances of “gmxapi.commandline.cli” are not by themselves conducive to DAG representations of data flow, so a helper function creates additional gmxapi primitives to provide a consistent set of named inputs and outputs. “gmxapi.commandline_operation()” generates a graph of gmxapi primitives around “cli” to translate input and output arguments into operation inputs and outputs, as specified by the user (Figure 3b).

In order to provide extensibility, arbitrary user code can be transformed into a gmxapi operation by decorating a Python function definition with “@function_wrapper” (Figure 3c). As an example, “commandline_operation” is written in this manner.

Looping and conditional commands are critical to adaptive ensemble simulation[25] and thus form a key part of the gmxapi repertoire. Conditional iteration takes the form of conventional gmxapi command syntax but relies on some metaprogramming under the hood. The “while_loop” command allows a graph to be dynamically extended as a chain of repeated operations, including fused operations constructed with the “subgraph” tool. A subgraph allows inputs and outputs to be expressed in terms of other gmxapi operations. When used with “while_loop”, internal subgraph state can be propagated from one iteration to the next to provide a consistent environment similar to a standard loop.

### The work graph

The work graph permits data flow topology to be represented independently from execution strategies or run time resource assignment details. An array of input sources provided to a command generates a corresponding set of tasks, such as for trajectory ensemble simulations. Because such ensembles may be coupled (and gmxapi currently does not have a mechanism to specify that an ensemble is uncoupled), arrays of simulations must currently be co-executed through the gmxapi mpi4py executor. Operations defined with “@function_wrapper”, including the other built-in operations, are assumed to be uncoupled, and are launched in a sequence determined by the DAG topology during recursive resolution of data dependencies, executed sequentially by the simple built-in gmxapi 0.1 executor.

The work graph enables straightforward dependency resolution: commands are specified to operate on abstracted “handles” or references to work inputs rather than requiring the fully instantiated objects, so chains of simulations can be expressed where one command depends on the outputs of a prior command. Each individual command is then ready to execute once its inputs have been fully resolved. Because operations and data flow are represented as a DAG, arbitrarily complex topologies can be expressed unambiguously without unexpected side-effects. Trajectories can be forked or extended without re-executing simulation segments or overwriting previous results.

### Execution and control flow

The gmxapi interface enables several levels of control logic, described below. Native Python logic (if/else statements, for loops, etc.) can be used with gmxapi operations. However, since gmxapi operations construct a work graph that is then executed, expressing conditionals within this work graph requires special constructs rather than standard Python operators. Gmxapi therefore provides logic such as “while_loop” as an efficient way to express adaptivity within a sequence of gmxapi operations. Python-level logic can be utilized by forcing work graph execution using the result() method, which causes explicit execution of all code required to produce the requested result. In contrast, gmxapi-level logic operates on Futures and is thus deferred until graph execution.

Gmxapi also provides low-level logic for simulation control and ensemble operators that can be used by third party code. For instance, gmxapi registers with the GROMACS “StopSignal” facility, enabling plugin code to stop a simulation based on external criteria, such as when a statistical estimator has converged. Additional examples of low-level operations include a “ReduceAll” to collect data across an in-flight ensemble. This operation can be used for adaptive updates across an ensemble of simulations, such as modifying biasing forces in restrained-ensemble simulations[26,27] or to update estimators and determine when a simulation ensemble should be terminated.

### Gmxapi as a means to extend MD code without source modification

Gmxapi establishes a facility[28] for providing MD extension code to GROMACS during launch. Binary objects are then loaded at runtime via the Python interpreter. This allows custom code to be executed during the MD integration loop with minimal overhead. Python facilitates plugin binding, but once the simulation launches, GROMACS and the extension code communicate directly via C++ interfaces, so there is no overhead added by Python. This extension code executes within the existing GROMACS parallelism framework, so it can take advantage of domain decomposition and MPI parallelism. Since this only requires a standard GROMACS installation with gmxapi, it permits flexible extension of the molecular dynamics code without modifying the main codebase or libraries.

Gmxapi does not prescribe a C/C++ bindings strategy. We used pybind11 for GROMACS bindings, but Python interaction with compiled code relies only on the Python C API (PyCapsule) and the public libgmxapi C++ interface.

## Results

Typical molecular dynamics workflows use a chain of tools to prepare simulation input, particularly command-line programs that take file locations as input and output arguments. Gmxapi is designed to permit connecting the output of any tool to the input for another tool with consistent Python syntax. Figure 4 illustrates a chain of GROMACS tools preparing simulation input, culminating with a call to “gmxapi.mdrun” to execute an array of simulations.

**Figure 4:**
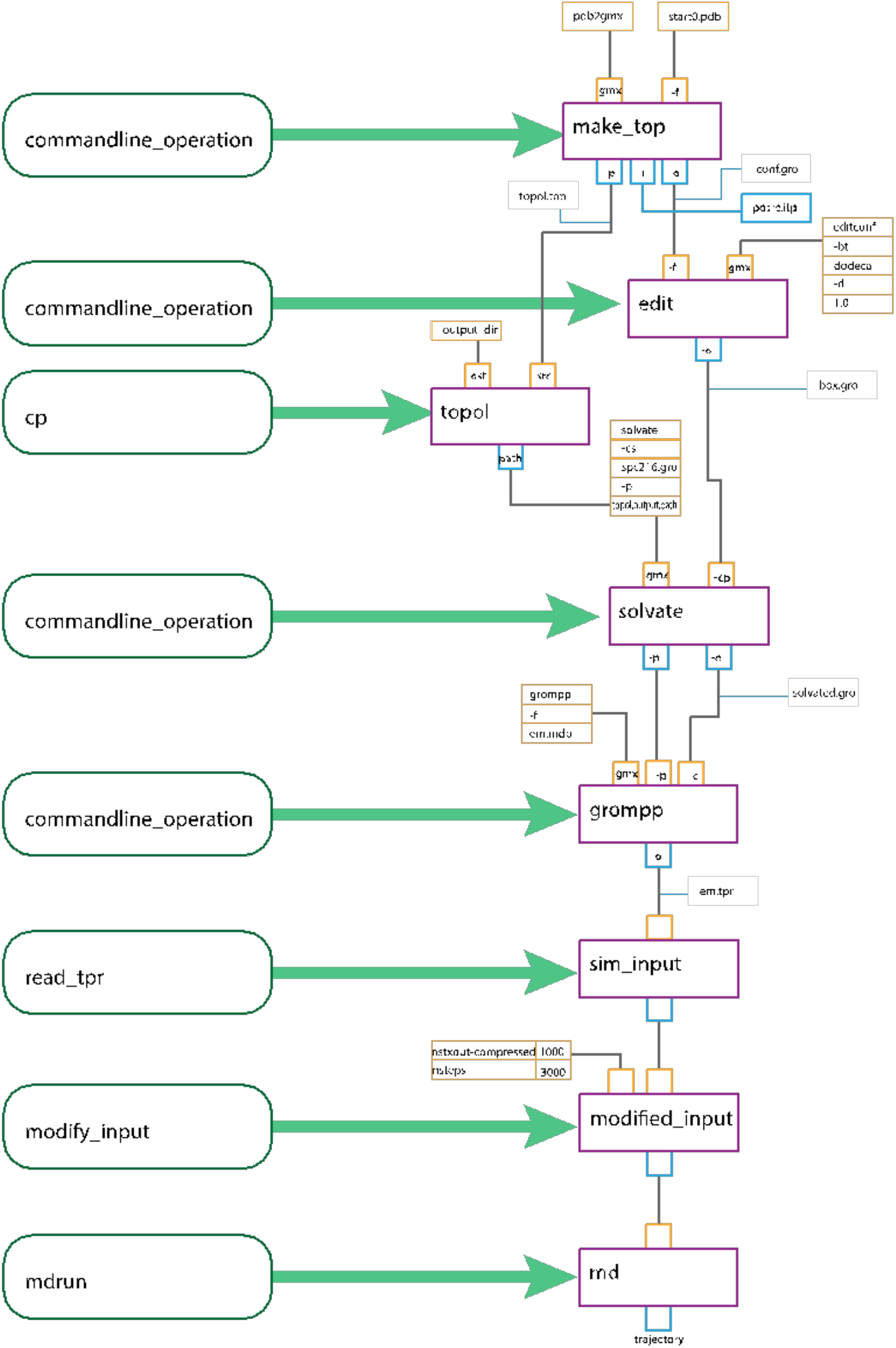
Detailed diagram of inputs, outputs, and operations in a chain of gmxapi operations. Operations are shown in green boxes, the corresponding nodes in the work graph in red boxes, inputs in orange boxes, and outputs in blue boxes. The diagram depicts a chain of GROMACS tools preparing simulation input, feeding into “gmxapi.mdrun” to run a batch of simulations.

To illustrate the application of gmxapi, we demonstrate refinement of an HIV gp41 conformational ensemble based on DEER spectroscopy data. We use a starting crystal structure for the BG505 SOSIP of HIV gp41[29] and previously reported DEER spectroscopy data of 5 different spin labels positions on this SOSIP[30]. The gp41 trimer was asymmetrically restrained (each of the three monomer-monomer distances was sampled separately), and the previously reported Bias-resampling ensemble refinement (BRER) algorithm for refining heterogeneous conformational ensembles[28] was applied. This involves a custom force plugin for GROMACS, and gmxapi was used to run an array of 250 refinement simulations, each randomly sampling target distance values from the experimental distribution for each measured spin-label pair. This array was executed in parallel on a supercomputing cluster, and accompanying example scripts demonstrate such a large-scale deployment. Further details are given in the Supplement. Full convergence of the refinement would require additional simulation sampling of the 15-dimensional experimental distribution, but even after 24 wall-clock hours of simulation a reasonable sampling of the experimental distributions was obtained (Figure 5). This application illustrates the ease of simulation ensemble management and application of custom biasing algorithms within GROMACS facilitated by gmxapi.

**Figure 5:**
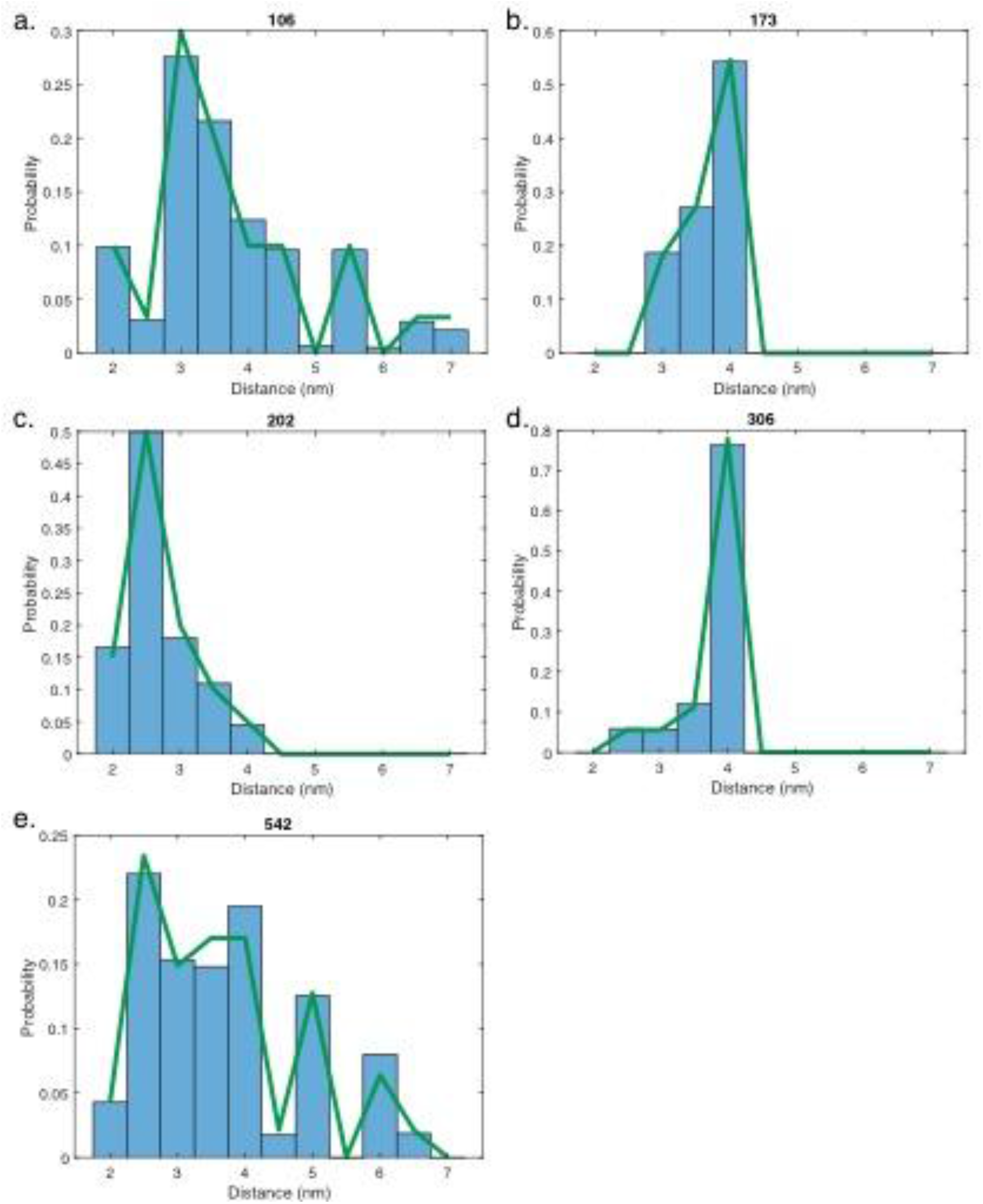
Residue-residue distance distributions in a simulated ensemble of HIV gp41. DEER spectroscopy was used to measure distance distributions between two monomers of the gp41 trimer. Thus, each residue designates a monomer-monomer residue pair. Panels a-e show plots for the 5 restrained residues in the HIV trimer: 106, 173, 202, 306, and 542 respectively. Discretized DEER distance distributions are plotted in green, and simulation results are plotted in blue bars. The simulation ensemble shows good convergence to the measured values.

## Discussion

Gmxapi provides a high-level Python interface for GROMACS with several key design features. It permits easy chaining of commands (GROMACS or third-party analysis tools) to create pipelines. Such pipelines can be parallel in nature, and gmxapi supports arrays or ensembles of simulations as first-class objects. Finally, gmxapi has a plugin interface that enables custom user code to be executed as part of molecular dynamics force calculations without modifying the GROMACS source but still benefitting from GROMACS native parallel decomposition. These features provide distinct advantages over shell-script-based workflows or simple Python wrappers for the GROMACS command-line interface.

Gmxapi differs from many molecular dynamics scripting APIs in that it embraces a data-flow programming paradigm while at the same time aiming for simplicity and ease of programming. This is analogous to the Keras high-level deep learning library [24]. A data-flow approach simplifies treatment of dependencies in complex parallel workflows and has been adopted by a number of high-performance computational tools.

Gmxapi scales well to high-performance clusters using MPI parallel interfaces. Current versions do not, however, provide advanced scheduler management along the lines of the prior Copernicus software [11-12] or the Radical Cybertools toolkit [16]. Interfaces that permit integration of gmxapi with more advanced schedulers is planned for future work. In comparison with Copernicus, gmxapi is a lighter-weight solution, integrating more closely with GROMACS and designed to facilitate much greater extensibility and ease of use but not including the advanced scheduler and client-server communication capabilities.

## Availability and Future Directions

The libgmxapi C++ interface was released as part of GROMACS 2019 release and has been part of a standard GROMACS installation since 2020. Gmxapi 0.2 shipped with GROMACS 2021; version 0.3, described here, is part of the GROMACS 2022 release and contains further enhancements for ease of use and installation. Continued development of gmxapi will expand the number of bound GROMACS API calls and reduce the need for legacy command-line support as well as generalize the plugin interface. At the programming level, one key future direction is exactly that—an increased use of the “Future” paradigm in Python. Python design patterns using Futures are becoming more widespread and standardized [14,31] as a way to refer to results that have not yet been calculated. Some additional refinement of the gmxapi Future protocol is needed to be fully compatible with native and third-party frameworks for concurrent or asynchronous program flow.

Code and data availability are specified below:

The gmxapi Python package is also maintained as part of the GROMACS repository at https://gitlab.com/gromacs/gromacs. It can be installed from the GROMACS source (https://gitlab.com/gromacs/gromacs/-/tree/master/python_packaging/src) or from https://pypi.org/project/gmxapi/ with “pip”, but needs to be told which GROMACS installation to use. The documentation at https://manual.gromacs.org/current/gmxapi provides details. During installation, the gmxapi Python package builds a C++ extension module against the GROMACS installation. Gmxapi tutorials are available from https://github.com/kassonlab/gmxapi-tutorials.

Custom molecular dynamics extension code is illustrated in a “sample_restraint” package: https://gitlab.com/gromacs/gromacs/-/tree/master/python_packaging/sample_restraint. BRER restraint potentials[28] are forked from this sample code and can be found at https://github.com/kassonlab/brer_plugin. Scripted BRER workflows are available at https://github.com/kassonlab/run_brer. The “run_brer” scripts were developed prior to gmxapi 0.1 and use internal logic to manage some data flow and to maintain workflow state. Both “run_brer” and “gmxapi” are being updated to illustrate that data flow and workflow state can be managed by the framework to allow simpler, more robust application code for applying new methods like BRER. Input data for the BRER example shown have been deposited at doi:10.5281/zenodo.5122931.

Gmxapi issues are tracked with the label “gmxapi” at https://gitlab.com/gromacs/gromacs/-/issues. Code contributions follow the GROMACS contribution procedure (https://manual.gromacs.org/current/dev-manual/contribute.html). However, gmxapi is intended to allow for maximal extensibility without requiring modification to the sources.

A discussion forum is available at https://gromacs.bioexcel.eu/tag/gmxapi.

## Acknowledgements

The authors thank Mark Abraham for many helpful discussions. This work was supported by a MolSSI Software Fellowship to M.E.I. and R01 GM115790 and OAC-1835780 to P.M.K. Computational resources on Frontera were provided under NSF LRAC MCB20006.

## Supporting Information contents

Supplementary Methods

Software Archive

## Notes

### Competing Interest Statement

The authors have declared no competing interest.

### Summary of Updates

Textual edits to improve clarity, new Figure 1.

