## Supplementary Information for "gmxapi: a GROMACS-native Python interface for molecular dynamics with ensemble and plugin support"

**Simulation Details.**

The crystal structure of the HIV gp41 BG505 SOSIP[1] was retrieved (PDB ID 4ZMJ) and prepared using CHARMM-GUI glycan reader[2], parameterizing using CHARMM36[3]. Glycans were truncated; this will likely affect the conformational equilibria of unrestrained simulations but was judged a reasonable initial approximation for refinement subject to experimental distance data. The SOSIP was solvated in 150 mM NaCl (with 75,410 water molecules) and energy minimized using GROMACS. After a brief equilibration, an initial run input file was prepared for BRER refinement. Simulations were performed with temperature maintained at 310K using the velocity-rescaling thermostat[4] and pressure at 1 bar using the Parrinello-Rahman barostat. A 1-fs timestep was used as per CHARMM forcefield recommendations, and van der Waals interactions were truncated at 1 nm. Particle Mesh Ewald[5] was used to treat long-range electrostatics.

Simulations were run on the Frontera supercomputer using 1 node per ensemble member for one wallclock day (336,000 core-hours).

### References

1. Do Kwon, Y.; Pancera, M.; Acharya, P.; Georgiev, I. S.; Crooks, E. T.; Gorman, J.; Joyce, M. G.; Guttman, M.; Ma, X.; Narpala, S.; Soto, C.; Terry, D. S.; Yang, Y.; Zhou, T.; Ahlsen, G.; Bailer, R. T.; Chambers, M.; Chuang, G.-Y.; Doria-Rose, N. A.; Druz, A.; Hallen, M. A.; Harned, A.; Kirys, T.; Louder, M. K.; O'Dell, S.; Ofek, G.; Osawa, K.; Prabhakaran, M.; Sastry, M.; Stewart-Jones, G. B. E.; Stuckey, J.; Thomas, P. V.; Tittley, T.; Williams, C.; Zhang, B.; Zhao, H.; Zhou, Z.; Donald, B. R.; Lee, L. K.; Zolla-Pazner, S.; Baxa, U.; Schön, A.; Freire, E.; Shapiro, L.; Lee, K. K.; Arthos, J.; Munro, J. B.; Blanchard, S. C.; Mothes, W.; Binley, J. M.; McDermott, A. B.; Mascola, J. R.; Kwong, P. D., Crystal structure, conformational fixation and entry-related interactions of mature ligand-free HIV-1 Env. *Nature Structural & Molecular Biology* **2015,** *22* (7), 522-531.

2. Jo, S.; Song, K. C.; Desaire, H.; MacKerell Jr, A. D.; Im, W., Glycan Reader: automated sugar identification and simulation preparation for carbohydrates and glycoproteins. *Journal of computational chemistry* **2011,** *32* (14), 3135-3141.

3. Best, R. B.; Zhu, X.; Shim, J.; Lopes, P. E.; Mittal, J.; Feig, M.; Mackerell, A. D., Jr., Optimization of the additive CHARMM all-atom protein force field targeting improved sampling of the backbone phi, psi and side-chain chi(1) and chi(2) dihedral angles. *J Chem Theory Comput* **2012,** *8* (9), 3257-3273.

4. Bussi, G.; Donadio, D.; Parrinello, M., Canonical sampling through velocity rescaling. *J Chem Phys* **2007,** *126* (1), 014101.

5. Darden, T.; York, D.; Pedersen, L., Particle Mesh Ewald - an N.Log(N) Method for Ewald Sums in Large Systems. *Journal of Chemical Physics* **1993,** *98* (12), 10089-10092.
